# Structure and stabilization of the antigenic glycoprotein building blocks of the New World mammarenavirus spike complex

**DOI:** 10.1101/2024.10.01.616071

**Authors:** Guido C. Paesen, Weng M. Ng, Geoff Sutton, Katie J. Doores, Thomas A. Bowden

**Affiliations:** Division of Structural Biology, Centre for Human Genetics, University of Oxford, Oxford, OX3 7BN, UK; Research Complex at Harwell, Harwell Science and Innovation Centre, Didcot, OX11 0FA, UK; Electron Bio-Imaging Centre, Diamond Light Source, Didcot, OX11 0DE, UK; Kings College London, Department of Infectious Diseases, 2nd Floor, Borough Wing, Guy’s Hospital, Great Maze Pond, London, SE1 9RT, UK

**Author notes:** Corresponding authors: Guido Paesen, Email:* *Thomas A. Bowden, Email:.

**Keywords:** arenavirus, glycoprotein, structure, rational immunogen design, virus-host interactions

## Abstract

The spillover of New World (NW) arenaviruses from rodent reservoirs into human populations poses a continued risk to human health. NW arenaviruses present a glycoprotein (GP) complex on the envelope surface of the virion, which orchestrates host-cell entry and is a key target of the immune response arising from infection and immunization. Each protomer of the trimeric GP is composed of a stable signal peptide (SSP), a GP1 attachment glycoprotein, and a GP2 fusion glycoprotein. To glean insights into the architecture of this key therapeutic target, we determined the crystal structures of NW GP1–GP2 heterodimeric complexes from Junín virus (JUNV) and Machupo virus (MACV). Due to the metastability of the interaction between GP1 and GP2, structural elucidation required the introduction of a disulfide bond at the GP1–GP2 complex interface, but no other stabilizing modifications were required. While the overall assembly of NW GP1–GP2 is conserved with that presented by Old World (OW) arenaviruses, including Lassa virus (LASV) and lymphocytic choriomeningitis virus (LCMV), NW GP1–GP2 complexes are structurally distinct. Indeed, we note that when compared to the OW GP1–GP2 complex, the globular portion of NW GP1 undergoes limited structural alterations upon detachment from its cognate GP2. We further demonstrate that our engineered GP1–GP2 heterodimers are antigenically relevant and recognized by neutralizing antibodies. These data provide insights into the distinct assemblies presented by NW and OW arenaviruses, as well as provide molecular-level blueprints that may guide vaccine development.

**Importance:** Although the emergence of New World (NW) hemorrhagic fever mammarenaviruses poses an unceasing threat to human health, there is a paucity of reagents capable of protecting against the transmission of these pathogens from their natural rodent reservoirs. This is, in part, attributed to our limited understanding of structure and function of the NW glycoprotein spike complex presented on the NW arenavirus surface. Here, we provide a detailed molecular-level description of how the two major components of this key therapeutic target assemble to form a key building block of the NW arenaviral spike complex. The insights gleaned from this work provide a framework for guiding the structure-based development of NW arenaviral vaccines.

## Introduction

Arenaviruses (family *Arenaviridae*, order *Bunyavirales*) comprise a group of genetically diverse, single-stranded, ambi-sense RNA viruses. Several mammalian-borne arenaviruses (genus *Mammarenavirus*) have repeatedly demonstrated the ability to spill over from rodent hosts and cause hemorrhagic fever or neurological disease in humans (1, 2). Mammarenaviruses are split into OW and NW lineages, and the NW lineage is further divided into four clades (A–D). The OW lineage includes the highly pathogenic Lassa virus (LASV), found in West Africa (3), and LCMV, which is prevalent across the globe (4). NW arenaviruses are endemic to the Americas and include the causative agents of Argentinian (Junín virus, JUNV) and Bolivian hemorrhagic fever (Machupo virus, MACV), both of which are clade B viruses (5). Apart from the live-attenuated Candid#1 strain of JUNV, which is licensed for use only in Argentina, no vaccines are available for human use against NW arenaviruses (6).

The arenavirus envelope surface is decorated with trimeric glycoprotein (GP) spikes, which are responsible for negotiating host-cell recognition and entry (7–10). Each protomer of the GP trimer consists of three non-covalently linked subcomponents (11): a stable signal peptide (SSP), a membrane-distal GP1 attachment glycoprotein, and a membrane-anchored GP2 glycoprotein, which are the result of proteolytic processing of a single SSP–GP1–GP2 chain in the endoplasmic reticulum. Signal Peptidase (SPase) cleaves the chain between the long, conserved SSP and the nascent GP1–GP2 segment, which is then processed by subtilisin kexin isoenzyme 1 (SKI-1) (12). In LASV, SKI-1 cleavage promotes spike assembly by increasing inter-protomer complementarity at the trimer interface (13). Moreover, the liberated, SKI-I site containing loop at the C-terminus of each GP1 (herein referred to as SKI-loop) helps to stabilize the trimer by forming bonds with a neighboring GP1–GP2 heterodimer, whilst apically exposed SKI-site residues are involved in receptor binding (14).

To facilitate endocytosis, GP1 interacts with host-cell receptors, including transferrin receptor 1 (TfR1; used by clade B and D NW viruses), α-dystroglycan (LASV, LCMV, clade C NW viruses) and neuropilin-2 (Lujo virus; LUJV) (15–21). Reflective of the diversity of receptors and permissive host species, GP1 exhibits considerable sequence variation, in contrast to the more conserved GP2 (22). Furthermore, OW LASV uses the endosomal receptor, Lysosome-Associated Membrane Protein 1 (LAMP1), which helps to facilitate membrane fusion (23), whilst OW LCMV targets the mucin region of CD164 (24). To date, no such intracellular receptors have been identified to be utilized by NW viruses during host-cell entry.

Following host-cell attachment and internalization, GP1 is expected to detach from the spike in acidic endosomes, enabling GP2 to enact its role as a class-I-fusion protein (10). GP2-mediated fusion occurs through insertion of bipartite fusion domains (25) into the endosomal membrane. Merging of the viral and endosomal membranes creates a fusion pore, allowing release of viral ribonucleoprotein complexes into the cytoplasm, where viral gene transcription and genome replication take place (5, 26).

A wide range of structural information is available for the OW virus GP. Both GP1/GP2 subcomponents and higher-order OW GP complexes have been resolved, alone and in complex with receptors and with Fab fragments of neutralizing antibodies (13, 14, 27–36). In contrast, structural information about NW GPs is limited to isolated GP1 and post-fusion GP2 subcomponents, either alone (32, 37, 38) or in complex with Fabs (39–42) or with human TfR1 (43). The absence of molecular-level insights into the higher-order NW GP complex may be attributed, in part, to the metastable nature of GP2 and its non-covalent association with GP1.

Although the NW GP complex is the primary target of the neutralizing antibody response arising from infection (10), little is known about how GP1 and GP2, the antigenic building blocks of this key vaccine target, interact. Here, we address this paucity of information through X-ray crystallographic determination of JUNV and MACV GP1–GP2 heterodimers in complex with the Fab fragments of neutralizing antibodies. Structural elucidation required the introduction of a disulfide bridge between GP1 and GP2, which stabilized and allowed production of the protein. Our data provide blueprints that will assist ongoing vaccine development efforts against NW arenaviruses.

## Results

### Expression and purification of NW GP1–GP2 heterodimers

Given the importance of NW GPs as vaccine targets, we sought to characterize the structural basis for how the interaction between GP1 and GP2 stabilizes this metastable complex. Focusing on GP from NW JUNV, we created a construct with an inter-subunit GP1–GP2 disulfide bond, further incorporating modifications utilized for the production of LASV GP1–GP2^e^ complexes (where ‘^e^’ denotes ectodomain) as a guide (13) (**Fig.1, Supplementary Fig. S1**). These modifications included replacement of the SKI-1 cleavage site with a furin cleavage site for improved processing efficiency, and the introduction of a proline substitution to prevent helices from forming or extending, as also utilized in SARS-CoV-2 spike (44). Recombinant protein production was performed in insect (*Spodoptera*) cells. Yields of purified protein for constructs with cysteines introduced at positions 88 and 329 (termed JUNV GP1^88-329^GP2^e^), and 88 and 328 (JUNV GP1^88-328^GP2^e^) resulted in the highest yields of purified protein (**Supplementary Figs. S1B** and **S1C**). Additionally, SDS-PAGE and size exclusion chromatography analysis revealed successful inter-chain disulfide-bond incorporation, furin cleavage, and the formation of the expected heterodimeric species (**Supplementary Fig. S2**). Introduction of inter-chain disulfide bonds at equivalent positions in other NW-virus GP1–GP2 constructs, including MACV (at positions 88 and 340, MACV GP1^88-340^GP2^e^), were similarly successful (**Supplementary Figs. S2** and **S3**). Further, production of OW LASV GP1–GP2 and LUJV GP1–GP2 complexes with analogous construct modifications also resulted in successful production of processed recombinant protein, albeit to relatively lower levels of expression (**Supplementary Fig. S3**).

**Figure 1.**
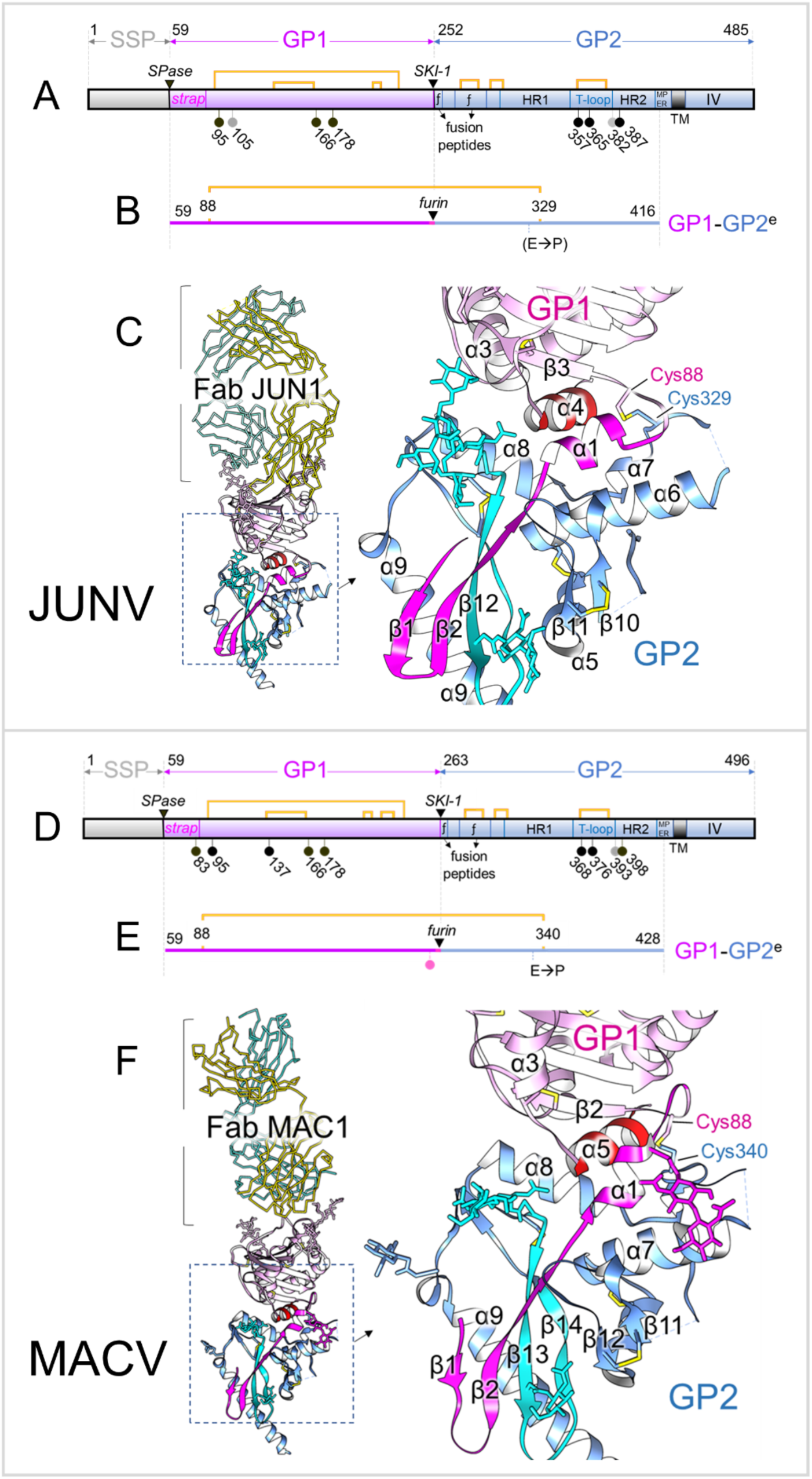
Structures of JUNV GP1–GP2 (top) and MACV GP1–GP2 (bottom). (**A**) Linear representation of the translation product of the JUNV spike gene, showing the SSP in grey, GP1 in violet and GP2 in pale blue. The black triangles indicate the SPase and SKI-1 cleavage sites, the pins represent N-linked glycosylation sites. Pins are colored black if the glycan was observed to be ordered and occupied, grey if not. The strap domain is indicated, as are the fusion peptides (ƒ), the Heptad Repeat regions (HR1 and HR2 (30)), the T-loop, the membrane-proximal external region (MPER), the transmembrane domain (TM) and the intra-virion (cytosolic) domain (IV). Disulfide bonds are indicated by yellow brackets. (**B)** Linear representation of the JUNV GP1^88-329^GP2^e^ construct. The signal peptide and twin-strep tag are not shown. The added disulfide bond is indicated by the yellow bracket, the introduced furin site by the black triangle. A proline (E321P) was also included, as incorporated in the study of LASV GP structure (13). The TM and IV regions were not included in the constructs. (**C**) Crystal structure of the JUNV GP1–GP2^e^ protein in complex with Fab fragments of neutralizing mAb JUN1. JUNV GP1–GP2 is shown in cartoon representation. The GP1 chain is in pink, except for the N-terminal strap (magenta) and the C-terminal α4-helix (red). The GP2 chain is colored blue, except for the T-loop region (cyan). The Fab from JUN1 is shown as ribbons, with the heavy chain in gold and the light chain in turquoise. Glycans are shown as sticks and colored according to the location of their sequon. Disulfide bonds are shown as yellow sticks. The insets show close-ups of the GP1–GP2 interface. The added cysteines forming the inter-chain disulfide bond are labelled. (**D** and **E**) Linear representation of the translation product of the MACV spike gene and of the MACV GP1^88-340^GP2^e^ construct, respectively, using the same color scheme and symbols as in panels *A* and *B*, but with E➔P denoting an E340P mutation. The mutation of Glu258 into a serine N-terminal of the furin-site in the MACV construct introduced an extra NXS glycosylation sequon (pink pin) to the ‘SKI-loop’. This loop is not seen in the crystal structure, so any bound glycans are not visible. (**F**) Crystal structure of the MACV GP1–GP2^e^ protein (cartoon) in complex with Fab fragments of neutralizing mAb MAC1 (ribbon). Colors and labelling are as in panel *C*.

### Structures of JUNV GP1^88-329^GP2^e^ and MACV GP1^88-340^GP2^e^

Furin-cleaved JUNV GP1^88-329^GP2^e^ and MACV GP1^88-340^GP2^e^ formed complexes with the Fab fragments of GP1-targeting neutralizing mAbs, JUN1 and MAC1, respectively (39), confirming the integrity of their folds. The complexes were crystallized and structurally elucidated to resolutions of 2.1 Å (JUNV–Fab JUN1) and 2.4 Å (MACV–Fab MAC1) (**Fig. 1**, **Supplementary Table 1**). As expected, the JUNV and MACV GP1–GP2^e^ structures are similar (**Fig. 1**, root-mean-square deviation (RMSD) of 1.9 Å), with differences localized predominantly to the 11 residue insertion in MACV GP1 (**Supplementary Fig. S4**), as described in previous structural studies (40, 45). JUNV and MACV GP1–GP2 are recognized by Fabs JUN1 and MAC1, respectively, at previously characterized epitopes located at their TfR1 recognition sites (39).

JUNV GP1 and MACV GP1 sit above the predominantly α-helical GP2s (**Fig. 1C** and **F**). Similar to their isolated structures, JUNV and MACV GP1 glycoproteins in GP2-bound forms consist of seven-stranded β-sheets (β3–9) interspersed with helical regions (**Supplementary Fig. S4**) (39, 41, 43). Overlay of GP2-bound JUNV and MACV GP1 onto their GP2 free states (PDB ID 7QU2 and 7QU1, respectively) resulted in low RMSDs (1.1 Å and 0.8 Å, respectively). This structural similarity is also reflected in structure-based classification analysis (**Fig. 2**), and indicates that unlike OW LASV GP1 (32, 46), the globular portion of NW GP1 shows only limited structural differences between the GP2-free and GP2-bound states (**Supplementary Fig. S5**).

**Figure 2.**
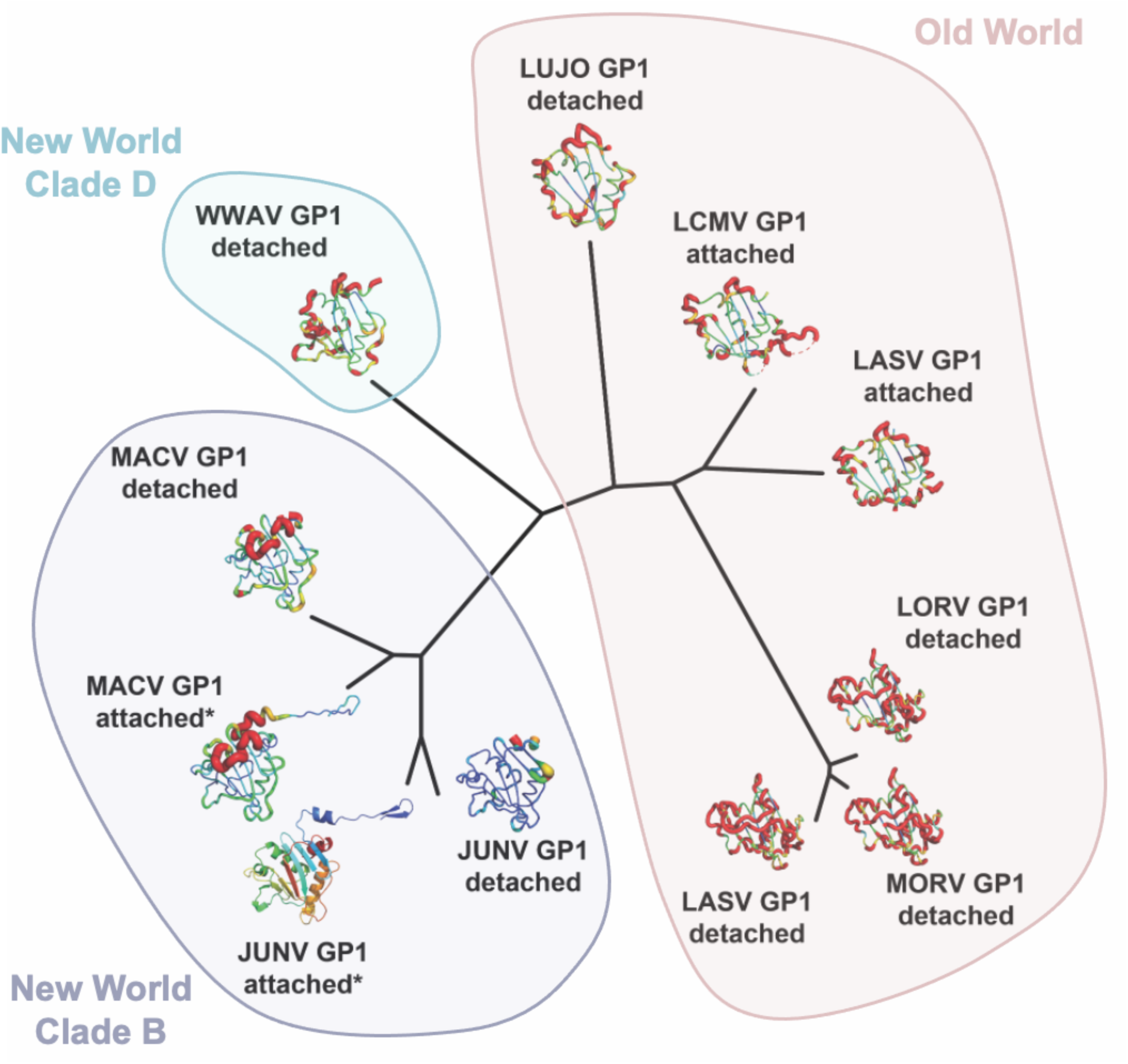
Structure-based classification of NW and OW arenaviral GP1 glycoproteins. Structure-based classification illustrates the similarity of NW GP1 glycoproteins in GP2 attached and detached states. A pairwise distance matrix was calculated with Structural Homology Program (SHP) (75, 76). Pairwise evolutionary distance matrices were used to generate un-rooted phylogenetic trees in PHYLIP (78). JUNV GP1, in the GP2 attached state, is shown in cartoon representation and colored as a rainbow from the N-terminus (blue) to C-terminus (red). Cartoon tube coloring from blue to orange, with increasing tube thickness, reflects increased structural distance from JUNV GP1 in the GP2-attached state upon overlay. Non-equivalent residues are colored red with exaggerated thickness.

In both JUNV GP1–GP2 and MACV GP1–GP2, the long and narrow, N-terminal ‘strap’ region (residues 59–89, **Fig. 3**) and the C-terminal α-helix (α4) of GP1 are visible, whereas they were previously not included in NW-GP1 constructs used for structural studies. However, no electron density corresponding to the loops at the C-termini of GP1, which are N-terminal to the cut furin cleavage sites, was observed, suggesting they are intrinsically flexible or may require the formation of higher-order trimers for stabilization.

**Figure 3.**
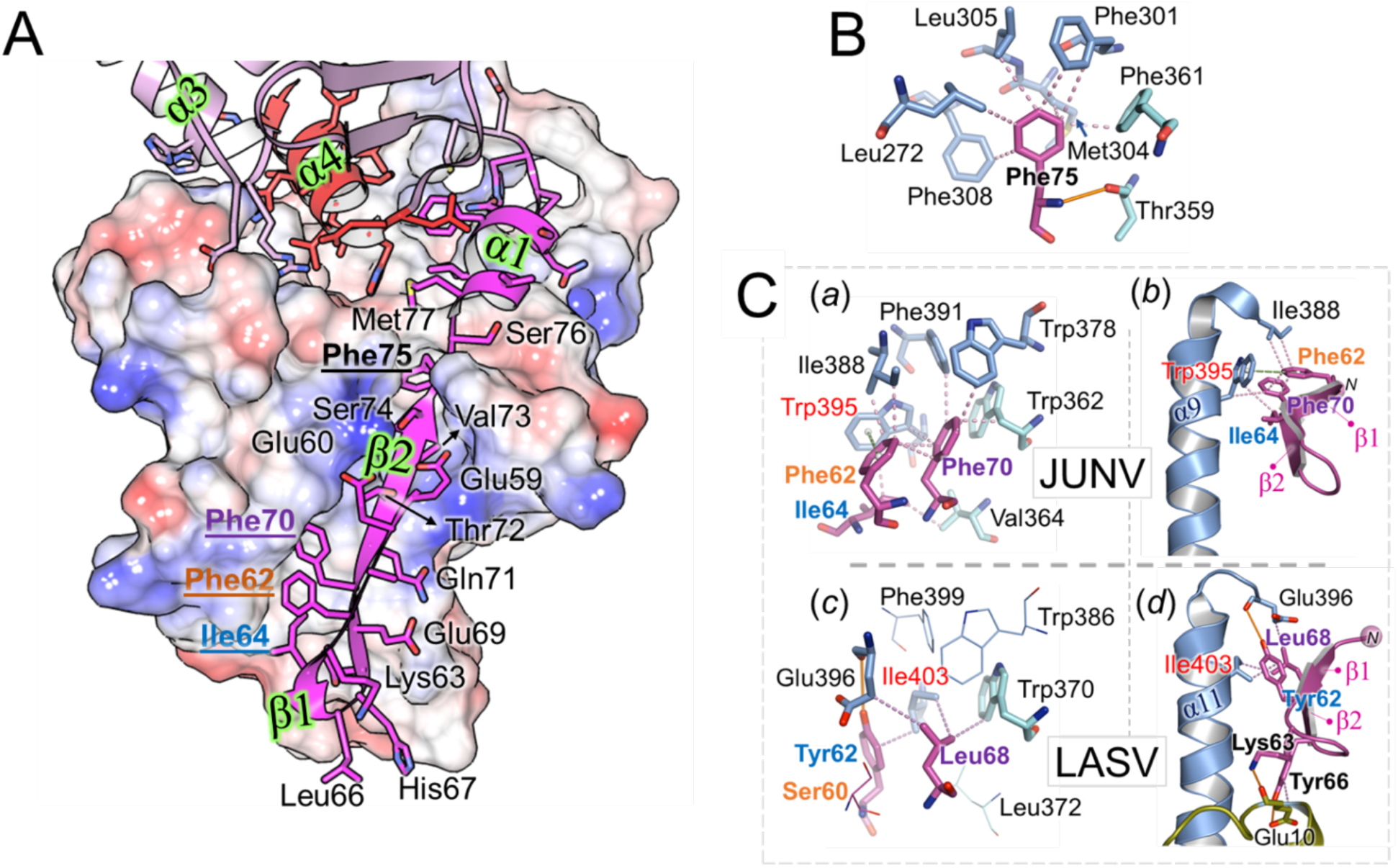
The N-terminal GP1 strap. (**A**) Overview of interactions between the GP1 strap and GP2. Given that the straps of JUNV and MACV are similar, the strap of JUNV was chosen for comparison with LASV. The GP1 strap is shown as a magenta cartoon, with side chains shown as sticks. GP2 is shown in surface representation, with basic and acidic patches colored blue and red, respectively. Residues occupying the long groove on the GP2 surface are labelled with bold letter types for Phe75 (in black), Phe70 (purple), Phe62 (orange), and Ile64 (blue). Labels for the helices and strands at the GP1–GP2 interface are highlighted in green. (**B**) Binding of Phe75 of the strap to a conserved, hydrophobic pocket in GP2. Strap residues are shown in magenta, GP2 residues in light blue or, if they belong to the T-loop, in cyan. Hydrogen bonds are shown as orange lines, hydrophobic bonds as violet dashed lines. GP2 residues forming the pocket are conserved in NW and OW viruses, but Phe75 of the strap is frequently replaced by a leucine, isoleucine, or valine (see **Supplementary Fig. S6**). (**C**) Comparison of the interactions at the N-terminal tip of the strap between the JUNV (*a, b*) and LASV (*c, d*) structures. In JUNV, Phe70 occupies a rather shallow, hydrophobic pocket, which consists of mostly aromatic GP2 residues (*a*). GP1 residues Phe62 and Ile64 join the pocket. Trp395 (labelled in red) plays a central role in the interaction, forming connections with Phe70 and Ile64, and π–stacking interactions with Phe62 (indicated by the green dashed line and spheres denoting the aromatic ring centers). Trp395 and Ile388 link the C-terminal helix of the ectodomain (α9; *b*) to the N-terminal β-strands of the strap (β1, β2; the N-terminus is denoted by ‘*N*’). Interactions in LASV (PDB 7PUY) at the level of Ile68 (equivalent to Phe70 of JUNV) are markedly different (*c*, *d*, and **Supplementary Fig. S6**). Ile403 (in red), which is strictly conserved in OW-viruses, takes the place of the much larger Trp395 (fully conserved in NW-viruses) on the C-terminal helix (α11, *d*). The smaller Ile403 partners with the aromatic Tyr62, which is also conserved in OW-viruses and replaces Ile64, the smaller, NW-specific partner of Trp395. Phe62 of JUNV, another residue that interacts with Trp395 and which is also conserved as a hydrophobic residue in NW-viruses, is replaced by Ser60 in LASV, which is not conserved in OW-viruses and does not contribute to the strap-GP2 interaction, as observed in PDB ID 7PUY (shown as lines). Similarly, Trp386 (Trp378 in JUNV), Phe399 (Phe391) and Leu372 (Val364) do not bind the N-terminus of the strap in LASV.

In our crystallized structures, MACV GP2 and JUNV GP2 exhibit the predominantly α-helical class-I pre-fusion fold presented by OW arenaviruses composed of N-terminal and internal fusion peptides, heptad repeat (HR) region 1 (subdivided into subcomponents a–d), T-loop, and HR region 2 (**Fig. 1**). Unlike the pronounced structural differences between OW and NW GP1s in the GP2 attached states (**Fig. 2**) the structures of OW and NW GP2 glycoproteins, in their pre-fusion GP1-bound states, are similar (∼1.9 Å RMSD). Structural deviation is highest at the C-terminal α-helix of the NW GP2 ectodomain (α9 in JUNV), where it assumes a similar orientation to that observed in trimeric LASV ectodomains (PDB ID 5VK2) but contrasts the membrane-anchored LASV GP spike (PDB ID 7PUY). Overall, the high level of structural conservation across the mammarenavirus GP2 is consistent with the evolutionarily conserved functionality of this subcomponent of the GP.

The engineered GP1–GP2 disulfide bonds in JUNV GP1^88-329^GP2^e^ and MACV GP1^88-340^GP2^e^ structures are occluded within their respective GP1–GP2 interfaces, where they are shielded by the loops they connect (α1–β2 and α6–α7 in JUNV) (**Fig. 1C**). The GP1 cysteine locates to the loop connecting the strap region with the central β-sheet, whilst the cognate cysteine in GP2 locates to HR1c, which is helical in reported LASV and LCMV GP1–GP2 structures (α9 of LASV, **Supplementary Fig. S4**). Interestingly, however, in the JUNV GP1^88-329^GP2^e^ and MACV GP1^88-340^GP2^e^ complex structures, only a small portion of the HR1c region is helical and JUNV Cys329 and MACV Cys340 reside within a loop that follows a helix-like trajectory.

In JUNV GP1–GP2, insect (*Spodoptera*) cell-derived glycosylation corresponding to paucimannose hybrid-type glycans was well ordered at three out of four N-linked glycosylation sequons (NXS/T, X≠P) of the GP1 subunit (Asn95, Asn166, and Asn178), and at two out of four sites (Asn357 and Asn365) on GP2. Glycosylation at all five and three out of four glycosylation sequons was observed in the GP1 and GP2 subunits of MACV GP1–GP2, respectively (**Fig. 1**). The extensive glycosylation of GP1 Asn95 in JUNV GP1–GP2 contrasts that observed in reported JUNV GP1 structures and may be attributed to stabilizing interactions imparted by GP2, or to the effect that construct boundaries and the expression system may have on glycan biosynthesis (**Supplementary Fig. S5**). Given the importance of glycans in epitope shielding of the arenavirus surface (47, 48), occupancy of N-linked sites constitutes an important consideration in the design of immunogens capable of accurately representing the antigenic GP.

### NW GP1 N-terminal strap interacts extensively with the GP2

The GP1–GP2 interfaces of JUNV and MACV structures occlude ∼2,250 Å^2^ and ∼2,170 Å^2^ of surface area, respectively (49) and are stabilized by extensive hydrophobic interactions, and 7 and 8 hydrogen bonds, respectively. This level of occlusion is similar to that observed in OW GP1–GP2 interactions (∼2,350 Å^2^ and 2,460 Å^2^ for LASV, PDB ID 7PUY, and LCMV, PDB ID 8DMI, respectively). The N-terminal strap, which consists of two β-strands (β1–2), a small α-helix (α1), and a C-terminal loop region preceding the central β-sheet in GP1 (**Figs. 1** and **3, Supplementary Fig. S6**), contributes the bulk of interactions with GP2, both in the JUNV (∼1,420 Å^2^ occluded surface) and MACV (∼1,380 Å^2^) structure.

The N-terminus of the strap region exhibits notable differences between OW and NW GPs in size, orientation, and how it connects to the C-terminal helix of GP2^e^ (**Fig. 3C**). In JUNV, this connection is largely defined by a tryptophan in the C-terminal (α9) helix (Trp395), which is strictly conserved amongst NW-viruses. In OW-viruses, Trp395 is invariably replaced by an isoleucine (LASV GP2^Ile403^), rationalizing the differential binding employed by the strap compared to NW viruses (**Fig. 3C**, **Supplementary Fig. S6**). This is also reflected in the strap sequences, where many of the GP2-interacting residues in JUNV and MACV are well conserved amongst other NW viruses, but not shared with OW viruses (**Supplementary Fig. S6**).

The main β-strand (β2) of the NW GP1 strap occupies a long and deep groove that traverses the outward-facing surface of GP2, joining the GP2 ‘T-loop’ β-strands (35) at the bottom of the groove to form a narrow but elongated antiparallel β-sheet. The β-turn (Leu66–His67) that connects β1 and β2 is located at a similar position to that in LASV GP, where it was shown to interact with the extracellular region of the SSP of the same protomer (14). In addition to main-chain hydrogen bonds between the β2 and β12/β13 strands, a series of conserved, hydrophobic interactions between GP1 and GP2 are observed, notably involving Phe70 and Phe75 (**Fig. 3B, C**).

The Phe75-containing loop preceding the α1-helix of the strap interacts at several points with the so called ‘internal’ segment of the fusion peptide (i-FPS (25)) via a series of main- and side-chain interactions (**Fig. 4A, B**). This arrangement suggests that dislodgment of the strap and the liberation of the fusion peptide in the endosomes are part of the same process. In line with this, the α1-helix and a short stretch of residues following the loop are linked to the N-terminal segment of the FPS, albeit indirectly, via α6 of GP2.

**Figure 4.**
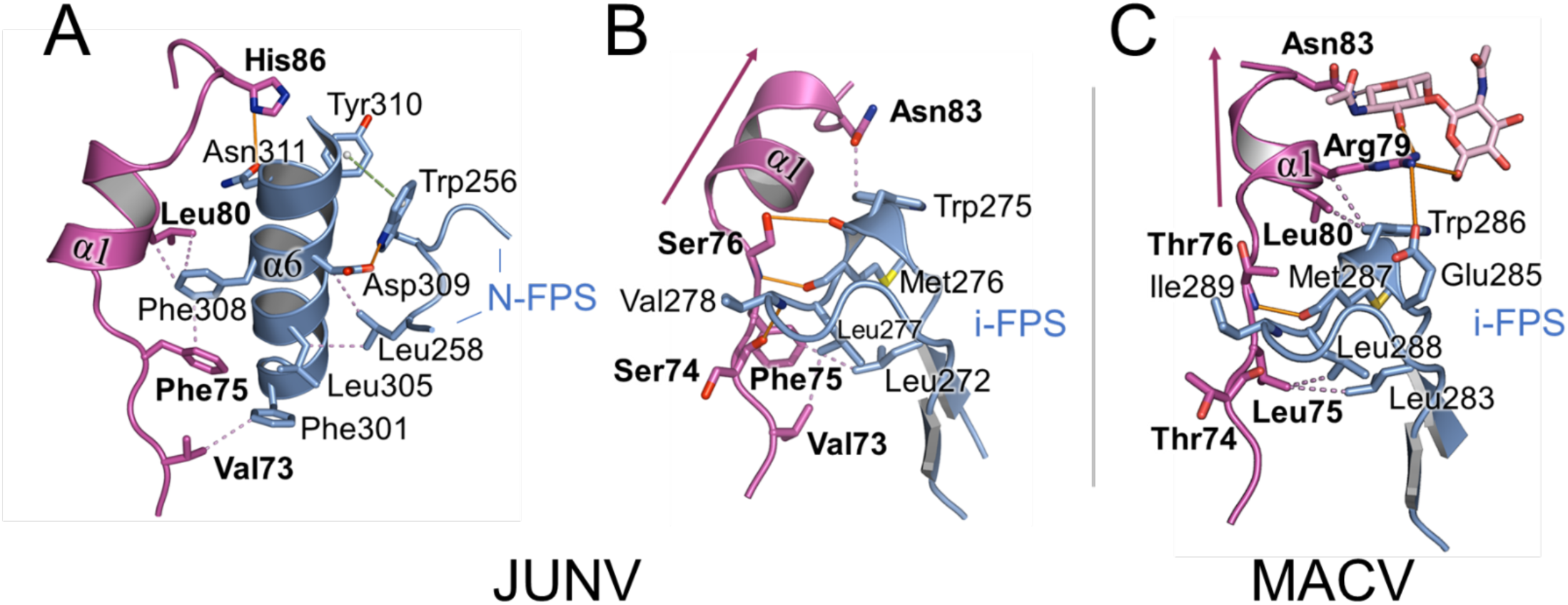
Interactions at the C-terminal region of the strap. **(A)** Interactions between the C-terminal part of the strap domain in JUNV and the α6 helix of GP2, which in turn interacts with the structurally ordered portion of the N-terminal fusion peptide segment (N-FPS). The strap domain is colored magenta, GP2 domains are colored blue. The side chains of selected residues and main chain atoms that form inter-chain hydrogen bonds are shown as sticks. Strap residues are labelled in bold. Hydrogen bonds are shown as solid orange lines, hydrophobic interactions as violet dotted lines, and π–stacking interactions as green dashed line with spheres denoting the aromatic ring centers. (**B**) Direct interactions between the C-terminal part of the strap in JUNV and the internal fusion peptide segment (i-FPS). (**C**) Interactions between the C-terminal part of the strap and the i-FPS in MACV. The purple arrow indicates the orientation of α-helix 1, which contrasts that observed in JUNV GP1. Instead of Asn83 interacting with i-FPS, as observed in JUNV GP1–GP2, Asn83 in MACV GP1–GP2 is distal from the fusion peptide and glycosylated. N-linked glycosylation is also found at the corresponding position in the GP1s of LASV (Asn79) and LCMV (Asn85), at a conserved NXS/T sequon at the C-terminus of η1 (**Supplementary Fig. S6**) and is similarly directed away from the fusion peptide.

The α1 helix (77–82) assumes different orientations in the JUNV and MACV GP1–GP2 complex structures (**Fig. 4**). Additionally, and possibly related to this observation, the neighboring Asn83 in MACV GP1 is glycosylated and directed away from the GP2-resident fusion peptide (**Fig. 4C**). Asn83 is part of a four-residue extension of the α1–β3 loop that is found in a small subset of clade B NW viruses, including JUNV, MACV, Tacaribe virus (TCRV), Tietê virus (50) and Ocozocoautla de Espinosa virus (OCEV) **(Supplementary Fig. S6)** (51). Although Asn83 is conserved amongst these viruses, only in MACV, TCRV and Tietê virus is it part of an NXS/T (where X≠P) N-linked glycosylation sequon. Interestingly, however, an NXS/T motif is commonly found at a similar location in OW viruses, C-terminal of the 3_10_-helix replacing MACV GP1 α1 (**Supplementary Fig. S6**). Similar to Asn83 in MACV, the corresponding Asn-residues in LASV (Asn79, PDB ID 5VK2) and LCMV (Asn85, PDB ID 8DMI) were observed to be glycosylated and directed away from the fusion peptide, indicative of a conserved structural feature across these distant arenaviral glycoproteins.

### The NW GP1–GP2 interface at the α3- and α4-helices

Both α3- and α4-helices of GP1 contribute to the NW GP1–GP2 interface (**Fig. 5**). For example, inter-subunit interactions occur between GP1 α3 and residues at and proximal to α8 of GP2. In JUNV GP1–GP2, this includes notable hydrogen bonds between Arg201 and the side chain and N-acetylglucosamine moieties of Asn357 (**Fig. 5D**). The α4-helix of the C-terminus of GP1 is located centrally in the GP1–GP2 interfaces. In addition to forming intra-subunit interactions with α1 of the strap, α4 is strongly embedded in the globular part of GP1 (**Fig. 5A–C**). Facing GP2, α4 wedges between Heptad Repeat (HR) regions 1c (α7) and 1d (α8) (30), which may sterically impede the formation of the continuous α-helix observed in the post-fusion conformation of GP2 (**Fig. 5B**).

**Figure 5.**
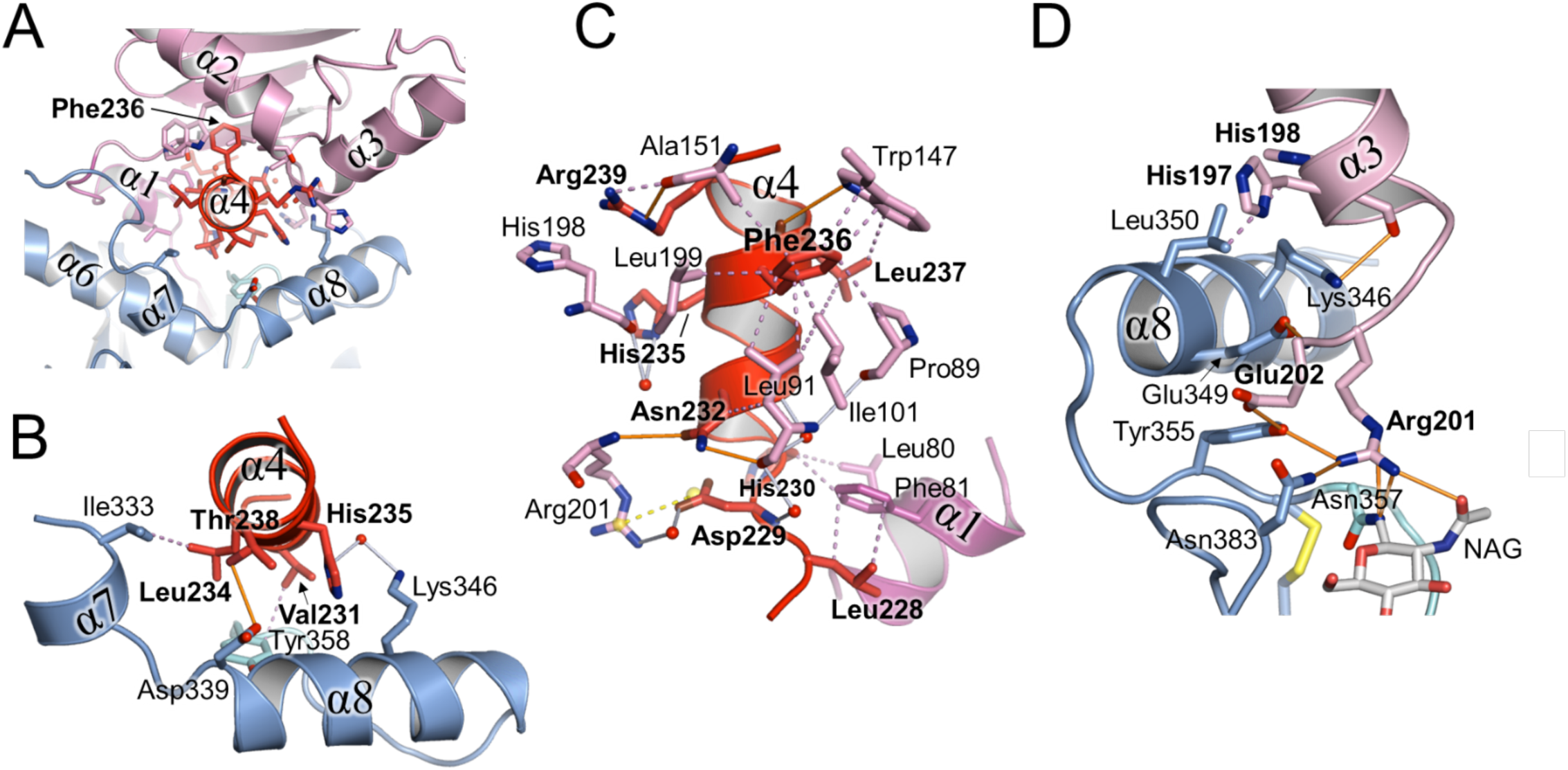
α-helix 4 of the GP1 is centrally located in the GP1–GP2 interface. (A) Overview, in cartoon representation, of the interaction between the α-helix 4 (colored red) of the JUNV GP1 (colored pink, except for α4) and JUNV GP2 (colored blue, except for the T-loop; cyan). The α4-helix is centrally positioned at the GP1–GP2 interface and embedded within the JUNV GP1 core structure, in part, through hydrophobic interactions with Phe236. (B) Close-up of the interface between α-helix 4 and GP2, which includes interactions between T-loop residue Tyr358 and α-helices 7 and 8. Hydrogen bonds are shown as orange lines, hydrophobic interactions as violet, dotted lines, and potential bonds via waters as white lines. (C) Interactions between α4 and the rest of the GP1 subunit. Phe236 forms part of a hydrophobic cluster comprised of Ala151, Trp144, Pro89, Ile101, Le91 and Leu199. Yellow dotted lines represent salt bridges. Interactions with α1 of the N-terminal strap are also shown. (D) Interactions between α3 of JUNV GP1 with GP2. Arg201 of GP1 forms interactions with residues adjacent to the disulfide bridge bordering the T-loop region (Cys356–Cys377; yellow sticks).

### The elongated morphology of JUNV and MACV GP1–GP2s

As described above, the conserved GP2^e^ subunits of JUNV and MACV share a high level of structural similarity with those of reported LASV structures (PDB codes 7PUY and 5VK2). However, upon superposition of the GP2 cores, we observe differences in the positions of the GP1s, relative to their cognate GP2s. Whilst maintaining the same orientation, the central β-sheets of the JUNV and MACV GP1s are at a greater distance from the GP2s than in LASV and LCMV GP1–GP2 structures (**Fig. 6A**). The relative ‘upward’ shift of the NW GP1, with respect to OW GP1, may be attributed, in part, to a four-residue extension of the loop following the α1 helix (**Supplementary Fig. S6**). This loop tethers the β-sheet to the GP2-embedded strap region. Given that only a small subset of clade B arenaviruses encode this loop extension, it is unlikely that the shift is a common feature among NW viruses (**Supplementary Fig. S6**). The upward shift, combined with the enlargement of α3 by the η2 helix, the addition of a strand (β6) to the sheet, and the protrusion of the β5–β6 and β7–β8 connecting loops from the apex of the GP1 fold, result in the observed elongated morphology of JUNV and MACV GP1–GP2 complexes, with respect to LASV GP1–GP2 (**Fig. 6A**). Accordingly, the distance between the membrane-proximal β1-β2 turn and the most apical GP1 residue is 80 Å in JUNV, compared to 70 Å in LASV. The extra β6-strand of GP1 may be a conserved characteristic of NW arenaviruses, where the loop containing it is generally much longer than in OW viruses (**Supplementary Fig. S4**).

**Figure 6.**
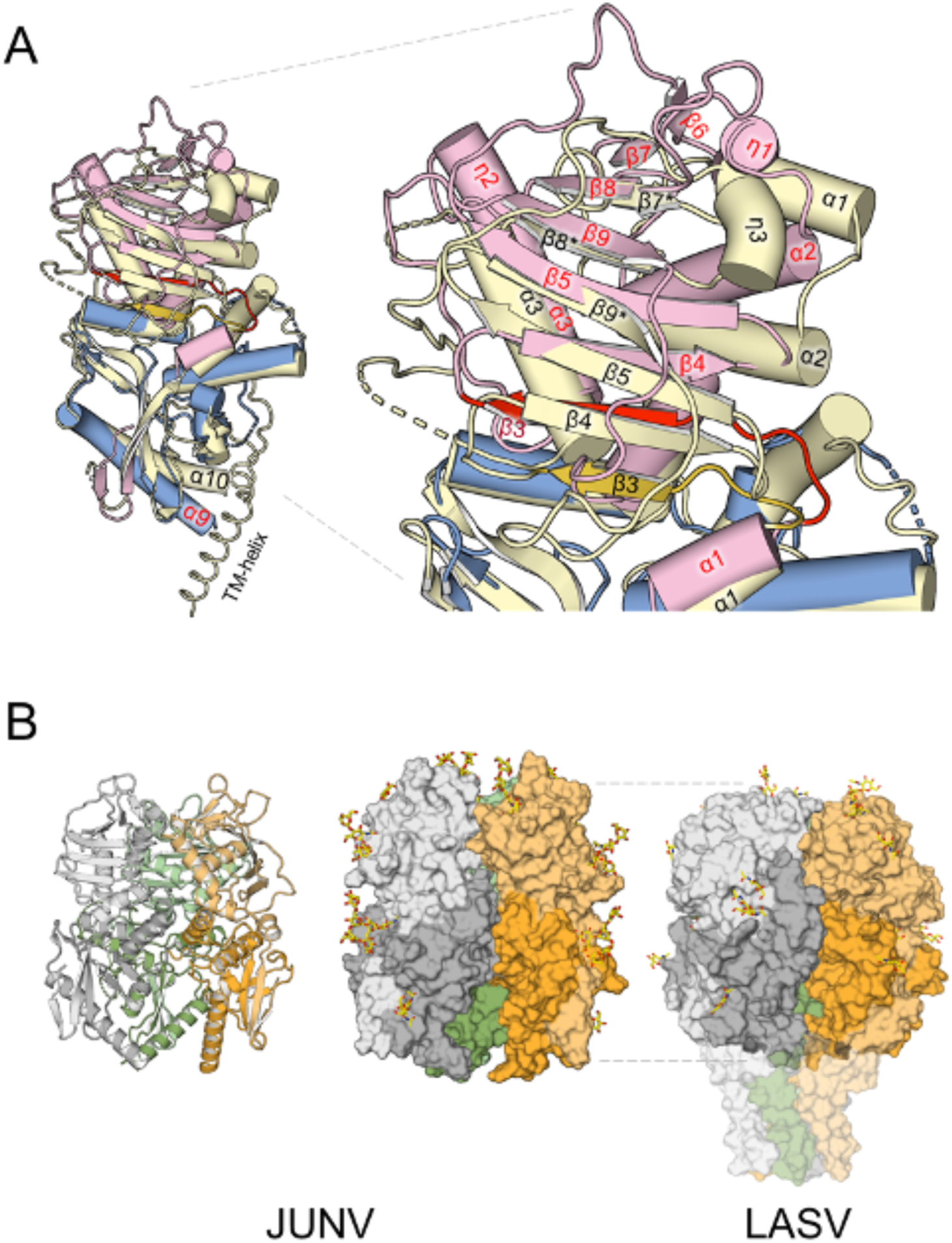
Comparison of GP1–GP2 heterodimers and higher-order trimeric assemblies. (**A**) The JUNV GP1^88-329^GP2^e^ structure, with GP1 (pink) and GP2 (blue), superposed onto LASV (7PUY) GP1-GP2 (yellow). The labels for the structural elements are colored red for the JUNV structure and black for the LASV structure. The symbols α, β and η indicate α-helices, β-strands and 3_10_-helices, respectively. The highly conserved GP2 subunits align well, apart from the helices at the C-terminus of the ectodomains (α9 in JUNV, α10 in LASV), which bend in different directions, likely due to the presence of the transmembrane region (TM) in the LASV structure. In JUNV and MACV, the loop that connects α-helix 1 of the GP2 embedded strap domain to β-strand 3 at the ‘bottom’ of the β-sheet in GP1 is longer than that in LASV. Loop and β3-strand are shown in red in JUNV and gold in LASV. The longer loop allows the entire β-sheet to move ‘upwards’, *i.e*. further away from GP2 than in LASV. (**B**) To model the trimeric JUNV spike, three copies of the JUNV GP1^88-329^GP2^e^ heterodimer were superposed onto a trimeric LASV GP structure (PDB ID 7PUY; (14)). The copies fit well into the modelled spike without major clashes, however, some flexibility may be required at the α2–β7 loop, which may otherwise be too close to 3_10_-helix 1 of a neighboring GP1 subunit. The model is shown in cartoon- (left) and surface-representation (middle), with the heterodimers shown in grey, green and orange, using lighter shades for the GP1 subunits. Glycans are shown as yellow sticks. The LASV structure (right) is shown for comparison, using the same color scheme, with the transparent surface showing the TM region. The model suggests the JUNV spike is more elongated than LASV, due in part to the upward shift of the GP1-sheet.

In JUNV, η1 and α2 assume a different orientation from that of the corresponding helices in LASV, where they demarcate a groove that acts as a binding site for the SKI-loop of a neighboring GP1 (14). A similar, well-defined groove is not obvious in our JUNV or MACV GP1–GP2 structures (**Fig. 7A**). Binding of SKI-loops in the native trimers of these viruses may thus involve a different set of interactions and a different trajectory of the loops, as is plausible in our model of the trimeric NW GP (**Fig. 7B**). This variation reflects the low level of sequence conservation of the SKI-loop amongst mammarenaviruses, where only the P4 residue of the cleavage site (Arg248 in JUNV, Arg256 in LASV) is strictly conserved.

**Fig. 7.**
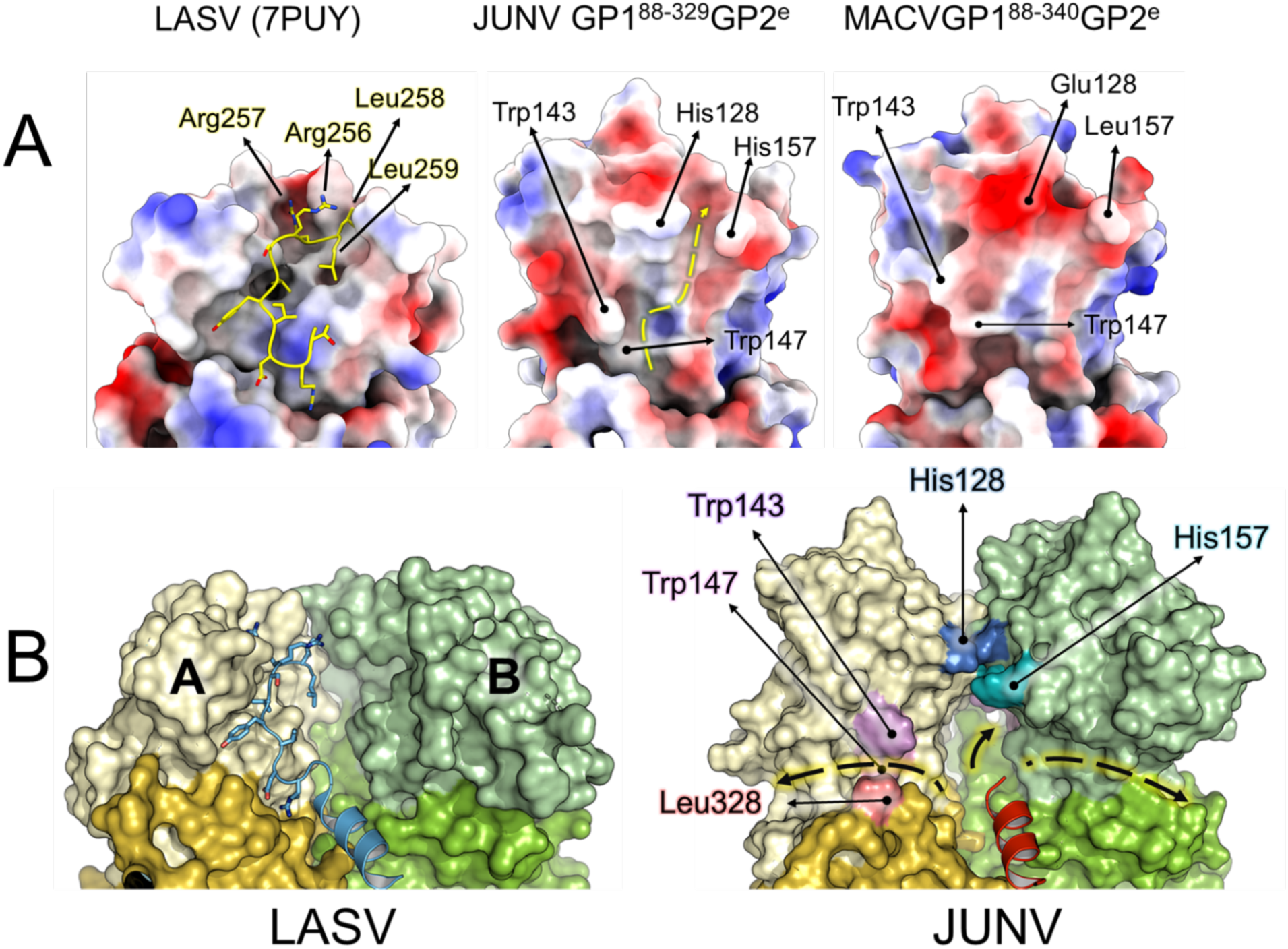
Mapping the putative trajectory of the SKI-loop. (**A**) The surfaces of LASV (PDB 7PUY), JUNV, and MACV GP1–GP2 heterodimers, colored from red (negative surface charge) to blue (positive), as seen from the interior of the trimer. The LASV GP1 surface presents a well-defined groove that can accommodate the C-terminal SKI-loop (*i.e.* the loop containing the SKI-1 recognition site following cleavage) protruding from a neighboring GP1 (yellow sticks) (14). In contrast, JUNV and MACV present a more negatively charged surface. Assuming LASV-like binding of the SKI-loop occurs in JUNV and MACV, the N-terminal end of the loop may be impeded by the side chains of Trp143 and Trp147, whilst the cleavage-site peptide could be sandwiched by the side chains of His128 and His157 (JUNV), or Glu128 and Leu157 (MACV). The yellow, dashed line indicates such putative trajectory in JUNV. (**B**) (left) Binding of the SKI-loop of protomer C (cartoon representation, blue) by protomer A (with GP1 in yellow, GP2 in gold), as it occurs in LASV GP (14). For clarity, the SKI-loops were deleted from protomers A and B. (right) Comparison with a trimeric JUNV GP model. The trajectory of the SKI-loop, as seen in LASV, is curtailed by the converging His128 and His157 side groups. However, the shift ‘upwards’ of GP1 (Fig. 6) and the change in orientation of α-helix 2, compared to LASV, may create space at the interface with GP2, between the protomers. This suggests alternative routes for the SKI-loops, as indicated by the arrows highlighted in yellow.

## Discussion

Our inability to protect against the unpredictable emergence of New World arenaviruses constitutes a substantial risk to human health and economy (52). Indeed, although Candid#1 protects against JUNV (6), the use of this vaccine is limited and there is a paucity of therapeutics capable of protecting against this and other zoonotic New World arenaviruses.

Neutralizing antibodies arising from infection are predominantly elicited against the NW GP (7, 39, 48). To better understand this key antiviral and vaccine target, NW GP1 and GP2 fragments have been subjected to high resolution structural studies, alone and in complex with receptors and Fab fragments of neutralizing antibodies (39–43). Despite the insights gleaned from these combined works, molecular-level detail of how these GP subcomponents assemble has remained elusive. Here, through elucidation of the GP1–GP2 heterodimeric architecture from the re-emerging and highly pathogenic JUNV and MACV, we refine our understanding of a key building block of the trimeric NW GP.

This work was dependent upon our ability to produce suitable amounts of metastable GP1–GP2 and required the incorporation of unique site-direct point mutations that help stabilize and maintain the pre-fusion conformation of the heterodimeric complex. Such an approach builds upon previous successes in introducing disulfide bond bridges to stabilize immunogens against other important pathogens, including LASV (13), foot-and-mouth disease virus (53), respiratory syncytial virus (54) and human metapneumovirus (55). Assessment of whether our NW GP1–GP2 constructs, when incorporated into immunogens, enhance the potency and breadth of the neutralizing immune response beyond that already demonstrated through immunization with GP1 fragments, will inform upon the development of next-generation vaccines.

Our incorporated disulfide bond links the loop preceding the central β-sheet of GP1 with the HR1c region of GP2. This linkage is at a different location from that used to stabilize OW virus GPs which connects the long α4–β8 loop of GP1 to the C-terminus of HR1d (13). Although we cannot preclude the possibility that our added disulfide bridge may impede proper formation of the HR1c helix, this region is occluded by the HR1b-loop and is therefore unlikely to markedly alter the antigenicity of the glycoprotein. Furthermore, our derived constructs are antigenically relevant and recognized by previously reported immunization-derived neutralizing monoclonal antibodies (39). Interestingly, we find that unlike the GP2-free state of OW LASV GP1, which when secreted has a putative role as an immune decoy (56), NW GP1 structures are highly similar between GP2-bound and GP2-free states.

Whether the JUNV GP1–GP2 and MACV GP1–GP2 structures described here are fully representative of the wider NW arenavirus lineage is uncertain. It is possible that the elongated shape of JUNV GP1–GP2 and MACV GP1–GP2, compared to OW virus GPs, may be more pronounced in the subset of clade B viruses that have a longer loop between the strap region and central β-sheet of the GP1. Indeed, other NW virus GPs may display a more compact fold that more closely resembles the morphology of OW arenavirus GPs. This hypothesis is especially plausible for the GPs of clade C viruses, which use α-dystroglycan as a cellular receptor and have SKI-1 site residues that are well conserved with OW arenaviruses, suggesting they may also interact with matriglycan groups (14).

Understanding the high-resolution structures of the glycoproteins presented by zoonotic arenaviruses constitutes a key step in the rational design of therapeutics that can increase our pandemic preparedness. While future efforts will no doubt focus on the engineering of homogeneously processed trimeric NW glycoproteins that present native-like glycosylation, this work provides much-needed molecular-level blueprints for the rational development of vaccines against newly and re-emerging NW arenaviruses.

## Materials and methods

### Strains

Accession codes and abbreviations are defined, as follows: ACO52428 (JUNV; *M. juninense*; Junín virus, XJ13 strain), AAN05425 (MACV; *M. machupoense*; Machupo virus, strain Carvallo), YP_089665 (SABV; *M. brazilense*; Sabiá virus); YP_001816782 (CHAPV; *M. chapareense*; Chapare virus), YP_001911113 (WWAV; *M. whitewaterense*, Whitewater Arroyo virus), YP_001649210 (OLVV; *M. oliverosense*; Oliveros virus), AAC32281 (PICHV, *M. caliense*; Pichindé virus), NP_694870 (LASV), P09991(LCMV), YP_002929490 (LUJV; *M.. Lujoense*; Lujo virus), AY129247 (GTOV; *M. guanoritaense*; Guanarito virus), UZO33083 (Tietê mammarenavirus), A0A023J4Z7 (LORV, *M. loeiense*, Loei river virus), AAN09948 (*M. tamiamiensi*: Tamiami virus), AAN32957 (*M. paranaense*; Paraná virus), AAG42529.1 (*M. allpahuayoense*; Allpahuayo virus), BAL03415 (*M. lunaense*; Luna virus), YP_009116790 (*M. gairoense*; Gairo virus), ABU94343 (Tonto creek virus), ABI97298 (Catarina virus), ABW96596 (Skinner Tank virus), YP_001649222 (*M. cupixiense*; Cupixi virus), YP_009553321 (Aporé virus), P31842 (TCRV; *M. tacaribeense*; Tacaribe virus), YP_010086246 (Xapuri virus), AFD98839 (OCEV; Ocozocoautla de Espinosa virus), YP_001649226 (*M. bearense*; Bear Canyon virus), AAT88084 (*M. piritalense*; Pirital virus), Q8B121 (*M.latinum*; Latino virus), P19240 (*M. mopeiaense*; Mopeia virus), YP_516226 (*M. praomyidis*; Mobala virus), AAN32967 (Amapari virus), QWQ58032 (Bitu virus), QLJ57221 (Dhati Welel virus), YP_010840421 (Kwanza virus), YP_010839773 (Alxa virus), AUF72664 (*M. wenzhouense*; Wenzhou virus), YP_001936019 (*M. flexalense*; Flexal virus), ADX32836 (Gbagroube virus), ADX32840 (Menekre virus), YP_009141005 (*M. okahandjaense*; Okahandja virus).

### Cloning

All enzymes were purchased from New England Biolabs. Synthetic DNAs were obtained from Geneart (ThermoFisher Scientific) and ligated into a modified pOPIN-vector (57, 58), putting them downstream of a p10 promotor and adding an enterokinase site and a Twin-Strep tag (59) to the C-terminus of the translation product. PCR-based mutagenesis was used to introduce stabilizing disulfide bonds and to alter the SKI-1 substrate peptide into a furin cleavage site. Whereas in the JUNV constructs, the replacement of the Arg–Ser–Leu–Lys SKI-1 site with an Arg–Arg–Arg–Arg peptide resulted in satisfactory furin cleavage, in the MACV GP1-GP2^e^, the residues flanking the SKI-1site needed to be additionally mutated into serines (Glu–Arg–Ser–Leu–Lys–Ala was changed into Ser–Arg–Arg–Lys–Arg–Ser). The sequences of all constructs were confirmed by Sanger sequencing (Eurofins). Plasmids were co-transfected with baculovirus DNA into *Spodoptera frugiperda* (*Sf*9) cells using Cellfectin II (Invitrogen) to generate recombinant baculovirus (58, 60). The quality of the baculovirus stocks was determined by monitoring cell lysis of infected cells with trypan blue (61).

### Expression and purification

Fab constructs were expressed in HEK293T cells as described before (39). Suspension cultures of baculovirus infected *Sf*9 cells in SF900II medium (Gibco; 27.5 °C, 120 rpm) were used for GP1-GP2^e^ production. Four days post-infection, the medium was supplemented with Tris pH 8.0 (to 10 mM) and EDTA (0.5 mM) and clarified (5000 x g, 45 min). BioLock solution (IBA; 2.4 ml/L) was added before the medium was passed over a column containing 0.5-ml Strep-Tactin XT resin (IBA). Resin-bound protein was washed with 100 mM Tris pH 8.0, 500 mM NaCl, 0.5 mM EDTA and eluted with this buffer supplemented with biotin (to 50 mM; Sigma). The eluate was concentrated using a 30 kDa molecular weight cut-off (MWCO) centrifugal filter device (Amicon, Millipore) and submitted to size-exclusion chromatography (SEC) over a Superdex 200 increase 10/300 GL column (Cytiva), using a 15 mM Tris (pH 8.0) running buffer containing 200 mM NaCl and 0.5 mM EDTA.

JUNV and MACV GP1–GP2^e^ containing fractions were further resolved over a 1-ml HiTRAP SP (HP) column (Cytiva) using a and a linear, 0-500 mM NaCl gradient (over 30 min, at a 1 ml/min flow rate) in a 30 mM Tris pH8.0 running buffer. Protein not retained by the SP column was applied to a 1-ml HiTrap Q column (Cytiva) for fractionation, using the same buffer and NaCl gradient.

Purified protein fractions were examined by SDS-PAGE over 4-20% Tris-Glycine gels (NuSep) and by western blotting, in the presence and absence of reducing agent (β-mercaptoethanol), to evaluate purity, disulfide bond formation and furin cleavage. For detection of protein on western blots, horseradish-peroxidase conjugated Strep-Tactin (IBA Lifesciences) was used in combination with luminescent Clarity Western ECL substrate (BioRad). Band intensities were measured using Invitrogen’s iBright analysis software.

### Crystallization, data collection, and structure determination

JUNV GP1–GP2^e^ and MACV GP1–GP2^e^ proteins were incubated for 3 h at room temperature with previously characterized, GP1-binding Fabs, termed JUN1 and MAC1, respectively (39). The resulting complexes were separated from excess Fab protein by SEC (as above). The samples were concentrated, and the buffer exchanged to 10 mM Tris pH 8.0, in 30-kDa MWCO centrifugal filter devices.

Crystals were obtained by vapor diffusion at 21 °C. Typically 100 nL of the GP1-GP2^e^-Fab complex (∼5mg/ml) was combined with 100 nL of reservoir solution in 96-well sitting drop plates (Greiner), and the mixture was allowed to equilibrate against 90 uL of reservoir solution (62). Crystals of the JUNV GP1-GP2 complex with JUN1 grew in 20% v/v 2-Propanol, 0.1 M Tris pH 8.0, 5% w/v PEG 8K, 6% 2-Methyl-2,4-pentanediol. The MACV GP1-GP2 complex with MAC1 crystallized in 90 mM Li2SO4, 90 mM Na2SO4, 90 mM K2SO4,12.5 % w/v PEG 4000, 20% w/v 1,2,6-Hexanetriol, 0.1 M Gly-Gly/AMPD pH 8.5. Diamond beamlines I03 and I04-1 (Harwell, UK) were used for diffraction data collection at 100K. Data processing employed the XIA2 suite (63) and structures were solved with the molecular replacement program PHASER (64), using previously determined coordinates of the JUNV and MACV GP1 subunits (39, 41), of the LCMV GP2 subunit (30) and of the Fabs (39). COOT (65) was used for model building, BUSTER (66) and Phenix (67) for refinement. Structure validation employed COOT and Molprobity (68). Data collection and refinement statistics are provided in **Supplementary Table 1**.

Figures were prepared using PyMOL Molecular Graphics System (Version 2.1) (69) and UCSF ChimeraX (49). ChimeraX was also used to calculate RMSD values. Stride (70) was used for secondary structure assignment. Fab residues were numbered using the Martin scheme of the Abnum numbering program (71).

Protein alignments were obtained using T-coffee (72) and visualised with ESPript 3.0 (73). Interactions were identified using PLIP (74).

### Structure-based classification analysis

For structural-based classification, the protein databank files of NW and OW GP1 monomers were prepared by removal of water molecules, ligands, and protein residues outside of the canonical fold. Structures were analyzed with the Structural Homology Program, SHP (75–77). Pairwise evolutionary distance matrices were used to generate un-rooted phylogenetic trees in PHYLIP (78).

### Data deposition

The atomic coordinates and structure factors for JUNV GP1^88-329^GP2^e^ and MACV GP1^88-340^GP2^e^ complexes with the Fab fragments of GP1-targeting neutralizing mAbs, JUN1 and MAC1, have been deposited in the Protein Data Bank with the accession codes 9GHJ and 9GHI, respectively.

## Acknowledgements.

The authors would like to thank Diamond Light Source (Harwell, UK) for beamtime (proposals 19946 and mx28534), and the I03 and I04-1 beamline staff for assistance with data collection. We thank Daniel Pinschewer and Mehmet Sahin for their collaboration, which helped to generate the mAbs, JUN1 and MAC1. This work was funded by Medical Research Council MR/L009528/1 and MR/S007555/1 (to T.A.B.), and MR/N002091/1 and MR/V031635/1 (to T.A.B. and K.J.D.). W.M.N is supported by the Wellcome Early Career Award 226997/Z/23/Z. The Centre for Human Genetics was supported by the Wellcome grant 203141/Z/16/Z.

